# The promoter mutation paucity as part of the dark matter of the cancer genome

**DOI:** 10.1101/2024.06.03.597231

**Authors:** Nicholas Abad, Irina Glas, Chen Hong, Annika Small, Yoann Pageaud, Ana Maia, Dieter Weichenhan, Christoph Plass, Barbara Hutter, Benedikt Brors, Cindy Körner, Lars Feuerbach

## Abstract

Cancer is a heterogeneous disease caused by genetic alterations. Computational analysis of cancer genomes led to the expansion of the catalog of driver mutations. While individual high-impact mutations have been discovered also in gene promoters, frequency-based approaches have only characterized a few novel candidates. To investigate the promoter mutation paucity in cancer, we developed the REMIND-Cancer workflow to predict activating promoter mutations *in silico*, irrespective of their recurrence frequency, and applied it to the PCAWG dataset. We positively validated 7 candidates by luciferase assay including mutations within the promoters of *ANKRD53* and *MYB*. Our analysis indicates that particular mutational signatures and necessary co-alterations constrain the creation and positive selection of functional promoter mutations. We conclude that activating promoter mutations are more frequent in the PCAWG dataset than previously observed, which has potential implications for personalized oncology.

## Introduction

Carcinogenesis is primarily driven by genetic mutations. Modern cancer therapy paradigms aim for the administration of drugs that molecularly target the weaknesses of individual tumors based on their mutational patterns. Such precision medicine programs report that around 30% of patients harbor actionable mutations, which can be addressed by targeted therapy^1,2^. Alas, the therapy successes reported for some cases are not universal. Explanations of the differential response are often missing^3–5^, but may be hidden in the long-tail of less frequent mutations of unknown significance. This includes uncharacterized coding and non-coding mutations, which affect transcribed as well as regulatory regions. A source for this variability of response exists but it is yet to be identified. To improve the rate of patients who might benefit from molecularly targeted drugs, a better understanding of this dark matter of the cancer genome is required.

Although cancer cells often accumulate hundreds to tens of thousands of alterations in their DNA in combinations that are unique for each individual patient^6^, only a small subset of these mutations, typically around four to five^6^, are considered pivotal driver events in the development of cancer. Single nucleotide variants (SNVs) are the most common type of genetic alteration, of which most are assumed to be non-functional passenger mutations, while others impact cell function to various degrees. Statistical analysis of characteristic footprints of positive selection are used to identify strong driver events^7^. Most of the known driver mutations directly impact the coding regions of genes, which make up only a small portion, approximately 1-2%, of the human genome^8^. In contrast, highly recurrent functional promoter SNVs (pSNVs) have been previously observed, most notably in regulatory sequences of two oncogenes, *TERT* and *CDC20*.

The hotspot mutations *TERT_C228T_* and *TERT_C250T_* ^9,10^ create transcription factor binding sites (TFBS) for ETS-family factors, which lead to an upregulation of the downstream *TERT* gene. *In vitro* luciferase assays were performed to quantify the impact of the mutations. The mutant promoter displayed 20% to 300% upregulation compared to the wild type allele varying based on the cell line and promoter fragment utilized^9–12^.

An alternative mechanism was described within the *CDC20* promoter. Rather than creating a TFBS, the *CDC20_G529A_* mutation was found to disrupt the binding of the well-conserved repressor ELK4 motif, which results in an upregulation of the downstream gene^13^. Subsequent *in vitro* validation showed that this pSNV led to a 50% increase in luciferase activity although this result was debated in another study^14^. Beyond these examples, little evidence for recurrent oncogenic promoter mutations is available.

In both cases, the primary feature for the identification of these functional pSNVs was mutational recurrence. A substantial number of studies ^8,15–18^ and tools, such as FunSeq2^19^, LARVA^20^, CompositeDriver^21^ and Mutpanning^22^ focus on this criterion. Other approaches prioritize features such as a pSNV’s functional impact bias or its DNA sequence context ^23–26^.

The remarkable paucity in functional pSNVs poses the question why such an apparently effective mechanism is scarcely represented among the known driver events. Genomic locations at which pSNVs can unfold driver potential may (a) be rarely mutated (mutation bias), (b) the selective advantages they offer may be insufficient to compete with coding driver mutations (selection bias), or (c) these non-coding events are elusive to detection by traditional recurrence-dependent detection algorithms (method bias).

Here, we build upon earlier approaches to identify pSNVs through cohort-based statistical analysis^8^. To also enable the discovery of singleton pSNVs (i.e. mutations that are uniquely observed in a single patient), we search for functional pSNVs that upregulate the associated gene. By expanding the search beyond reliance solely on apparent positive selection signals like recurrence, we widen the search space of precision oncology programs also to this part of the dark matter of the genome.

To this end, we created the *Regulatory Mutation Identification ‘N’ Description in Cancer* (REMIND-Cancer) workflow. It incorporates whole genome sequencing (WGS) data, RNA sequencing (RNA-Seq) data and additional annotation-based information. Candidate mutations are then prioritized, ranked, and manually curated using our data integration tool, *pSNV Hunter*, for functional validation *in vitro* by luciferase assays, thus following a filter-ranking-inspection-validation paradigm.

We applied the computational pipeline to 2,413 cancer samples across 43 distinct cancer cohorts from the Pan-Cancer Analysis of Whole Genomes (PCAWG) dataset^6^. Aiming for high specificity, the REMIND-Cancer pipeline identifies pSNVs that alter TFBS motifs in core promoters of upregulated genes (filter step). Subsequently, leveraging the specific features of each of these 19,250 mutations, a prioritization score was computed based on their genomic, transcriptomic, and annotation-based features (ranking step). Next, high-scoring candidates were subjected to a semi-automated curation process. The expression patterns of target genes and TFs matching mutated TFBS motifs were analyzed on the cohort level. Alignment artifacts were removed using our *DeepPileup* approach^5^. Footprints of convergent cancer evolution for implicated promoter-gene pairs were analyzed on a pan-cancer level using *GenomeTornadoPlot* (inspection step).

We tested 16 pSNVs *in vitro* by a luciferase reporter assay (validation step). Among these validated mutations, we demonstrated a statistically significant upregulation of their respective target genes within seven (44%) pSNVs, four of which are singletons. These results affirm the feasibility to discover novel functional pSNVs even if they only occur in a single patient sample. A more detailed analysis of our results yields evidence that mutation, selection, and method bias all contribute to the apparent pSNV paucity in cancer.

## Results

### Recurrence alone identifies only a few potentially functional promoter mutations

To identify putative functional pSNVs, the REMIND-Cancer workflow used a series of filters on data from the PCAWG project consisting of 2,413 total patients, 927 of which have matched WGS and RNA-Seq data across 24 cohorts. (Fig. 1a). The application of our workflow identified 366,373 SNVs in promoter regions (Fig. 1a). By applying the gene expression, TFBS motif and TF expression filters (see “Methodology”), the candidate set was reduced to 19,250 pSNVs, which occurred within 10,550 unique genes and 882 patient samples. These remaining pSNVs were then annotated according to mutational recurrence, tumor purity, allele frequency, inclusion of the matching gene in the Cancer Gene Census (CGC) database, and co-location with accessible chromatin. These features were used to compute an empirical prioritization score, which was then used for subsequent pSNV candidate ranking. Hereby, recurrent pSNVs were scored separately for each sample they occurred in.

**Fig. 1:**
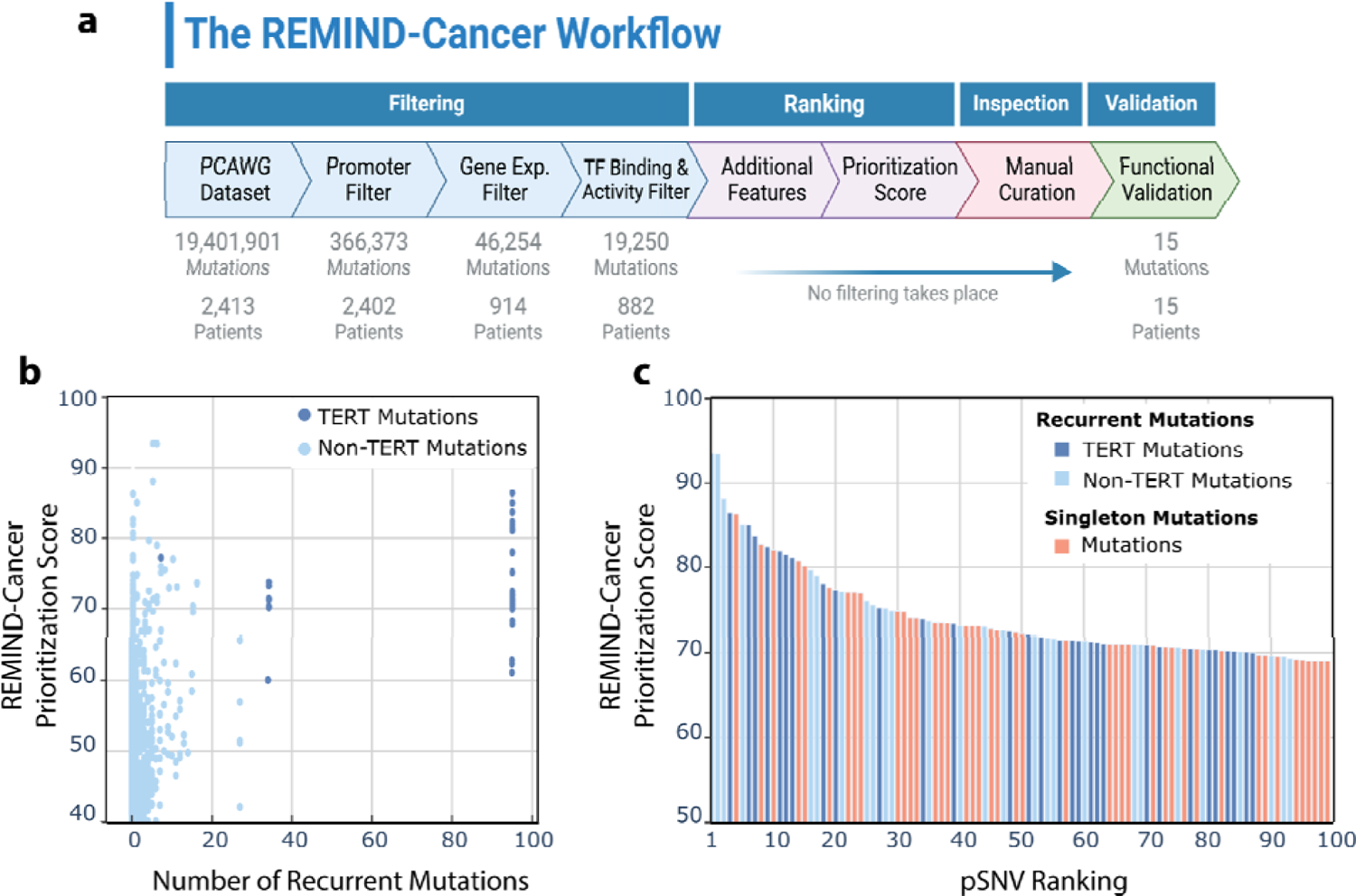
Characterization of the results of the REMIND-Cancer workflow. **(a)** An overview of the REMIND-Cancer workflow, which has four major components: filtering, ranking, inspection, and validation. Within each of these four components (i.e. filtering), the different steps (i.e. promoter filter) can be seen along with the number of mutations at the conclusion of this step (i.e. 366,37 3 mutations) and the number of patients that these mutations belong to (i.e. 2,40 2 patients)^27^ **(b)** For pSNVs that passed the pipeline, the sample-specific recurrence level (mutation frequency minus 1; x-axis) is plotted against its REMIND-Cancer prioritization score (y-axis). Both TERT promoter mutations are depicted in dark blue and the recurrent non-TERT mutations in light blue with TERT. **(c)** Distribution of the REMIND-Cancer Prioritization Score of the top 100 ranking pSNVs. Each bar represents a sample-specific pSNV and is colored by recurrence or singleton statu s and TERT association. With the current scoring parameters (Supplementary Fig. 1), the maximum score a pSNV can obtain is 117.

Overall, of the 96 *TERT_C228T_* and 35 *TERT_C250T_* pSNVs within PCAWG, 28 and 7 passed the gene expression filter, respectively. Due to a difference in normalized gene expression, each sample-specific pSNV receives a different prioritization score (Fig. 1b). Among the top 100 ranking pSNVs, 26 instances of *TERT_C228T_* and 7 instances of *TERT_C250T_* were represented (Fig. 1c). Moreover, our analysis also detected the previously-reported^13,14^ *CDC20_G529A_* pSNV in a single patient with elevated *CDC20* expression and a recurrence in 11 other samples, 2 of which had RNA-Seq data available.

It should be noted that the majority (n = 18,284 / 95%) of pSNVs passing the filtering steps were singletons. Beyond *TERT*, the only four other recurrent mutations within the top 100 ranking pSNVs with exactly two occurrences each were *ANKRD53_G529A_*, *SECISBP2 _G357A_*, and *RALY_C927T_* (Supplementary Data 1). In contrast, 46 of these top 100 mutations were singletons (Fig. 1c).

### The Recurrent *ANKRD53* Promoter Mutation Leads to an Increase in Promote r Activity through the Creation of a Binding Site for the NF-kB subunit p65

The *ANKRD53_G529A_* pSNV was recurrent in six patients, two of which had RNA-Seq data available. These two pSNVs were from different cohorts (bladder cancer BLCA-US and lung adenocarcinoma LUAD-US) and had the highest and third highest prioritization score, respectively (Supplementary Data 1). The prioritization score was derived from three major aspects: genomic, transcriptomic and annotation data, each comprising subcategories (see Online Methods). A detailed breakdown of the prioritization score for the highest-ranking *ANKRD53_G529A_* pSNV is illustrated in Fig. 2a in the form of a sunburst chart where the score exhibited high values for all three major categories.

**Fig. 2:**
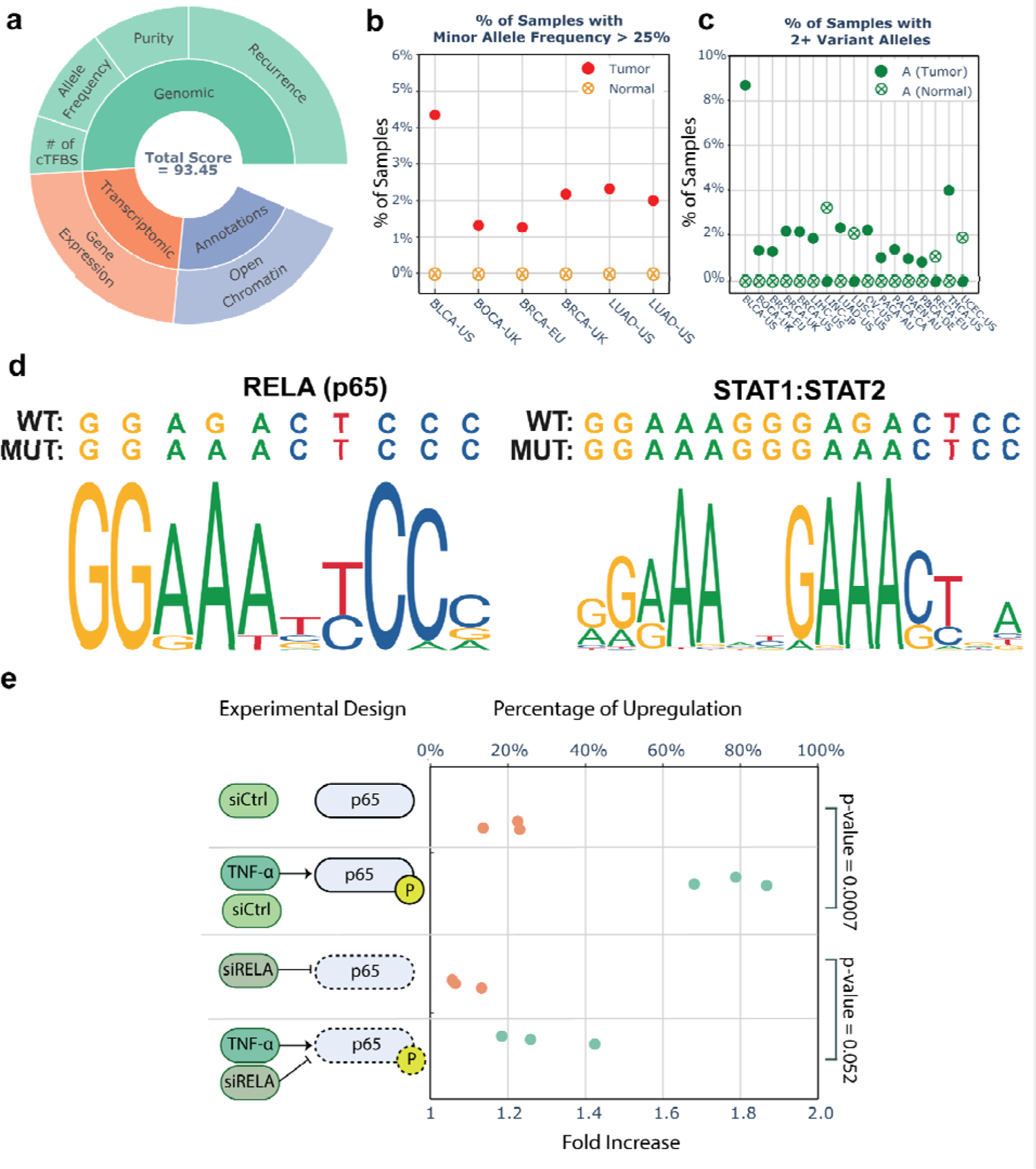
ANKRD53_G529A_ activates transcription in a NF-kB dependent manner **(a)** REMIND-Cancer scoring breakdown of the highest-ranking ANKRD53 pSNV within the BLCA-US cancer sample. The inner circle represents the three general scoring categories (genomic, transcriptomic and annotations) and the outer circle details the specific components of the inner circle. Here, the number of TFBSs that were predicted to be created by this pSNV is denoted by “# of cTFBS”. **(b)** A DeepPileup plot for quality control measures representing the percentage of samples per cohort in which the non-reference allele reaches a frequency greater than 25%. (c) A second DeepPileup plot representing the percentage of samples per cohort with two or more variant alleles at this position. (d) Logo plots (left: RELA (p65), right: STAT1:STAT2) of two TFBS’ which were predicted to be created by ANKRD53_G529A_ (e) The experimental design (left) and luciferase reporter assay results (right) of how the ANKRD53 promoter activity changes in response to the stimulation with TNF-α and expression reduction of p65 using siRELA. An upregulation of 0% represents no difference between the wildtype and mutant (i.e. fold increase of 1.0), whereas 100% represents activity doubling (i.e. fold increase of 2.0). The median of six technical replicates was determined for three independent biological replicates. The corresponding p-value is computed by a one-sided t-test.

Inspecting the *GenomeTornadoPlot and DeepPileup* diagrams for *ANKRD53_G529A_* indicated biological and technical validity of the pSNV. The *GenomeTornadoPlot,* which inspects the copy number alterations across the whole PCAWG cohort, displayed a mild enrichment of focal duplications overlapping with the gene (Supplementary Fig. 2).

*DeepPileup* at this position showed that only six PCAWG cohorts contained tumor samples with a minor allele frequency above 25% while all normal samples were negative (Fig. 2b). Counting samples with 2 or more reads carrying the adenine mutation showed 11 additional cohorts displaying a signal (8 tumor and 3 normal) as displayed in Fig. 2c. Although truly problematic positions show a higher background signal (Supplementary Fig. 3), we deemed functional validation to be necessary.

At the transcriptomic level, both samples harboring the *ANKRD53_G529A_* pSNV exhibited increased gene expression with a FPKM z-score of 1.42 and 1.76 in BLCA-US and LUAD-US, respectively. This G/C > A/T pSNV creates a highly-conserved base pair in each of the *de novo* TFBSs of the RELA and STAT1:STAT2 transcription factors as exemplified by the motif logo plots in Fig. 2d.

To measure the functional impact of the mutation, we established a validation workflow employing a luciferase reporter assay. Specifically, we designed a promoter fragment spanning 501 bp, encompassing the target pSNV (Supplementary Data 2). To account for co-regulatory elements, this fragment also incorporated regions with active histone marks observed in the UCSC genome browser^28^. The pSNV was predicted to create novel binding sites for the NF-kB subunit p65 (encoded by the *RELA* gene) and the STAT1:STAT2 heterodimer (Fig. 2d). The wildtype and mutant versions of this fragment were introduced into a reporter vector. Following transfection into HEK293FT cells, we observed a 20% increase in luciferase activity (Fig. 2e). Since NF-kB requires post-translational activation via its upstream pathway, we stimulated its activity using recombinant TNF-α under two conditions: downregulating the expression of the endogenous p65, the *RELA* gene product, with small interfering RNA (siRELA) or using a control siRNA (siCtrl). TNF-α stimulation increased the relative upregulation of promoter activity from 20% to 80% compared to the wildtype construct. This significant difference was reverted by siRNA-mediated p65 knockdown (Fig. 2e; Supplementary Data 3).

In summary, these results demonstrate the activating function of the *ANKRD53_G529A_* pSNV as well as one of the responsible transcription factors, thereby demonstrating the accuracy of the pipeline’s predictions in this specific case. It also showcases a potential additional layer of complexity to be considered for the study of functional promoter mutations, which derives from the activation and expression level of the transcription factors binding to mutation-induced *de novo* TFBS motifs.

### Five Additionally Identified Promoter Mutations Lead to a Significant Increase in Promoter Activity

Next, we utilized the prioritization score and our manual curation process to select and validate 14 additional mutations that we deemed to have potential functional significance. These selected mutations comprised both recurrent and singleton mutations, which were not exclusively associated with known cancer genes (Supplementary Data 4).

In total, five (36%) of the 14 additionally selected candidate mutations demonstrated a statistically significant increase in reporter activity (one-sided t-test), i.e. *CDC20_G529A_* (p-value = 0.016), *RALY_C927A_* (p-value = 0.0037), *PRKCB_C963T_* (p-value = 0.023), *EGR1_C049T_* (p-value = 0.016), and *TFEB_T989G_* (p-value = 0.002) (Fig 3a and 3b; Supplementary Fig. 4).

**Fig. 3:**
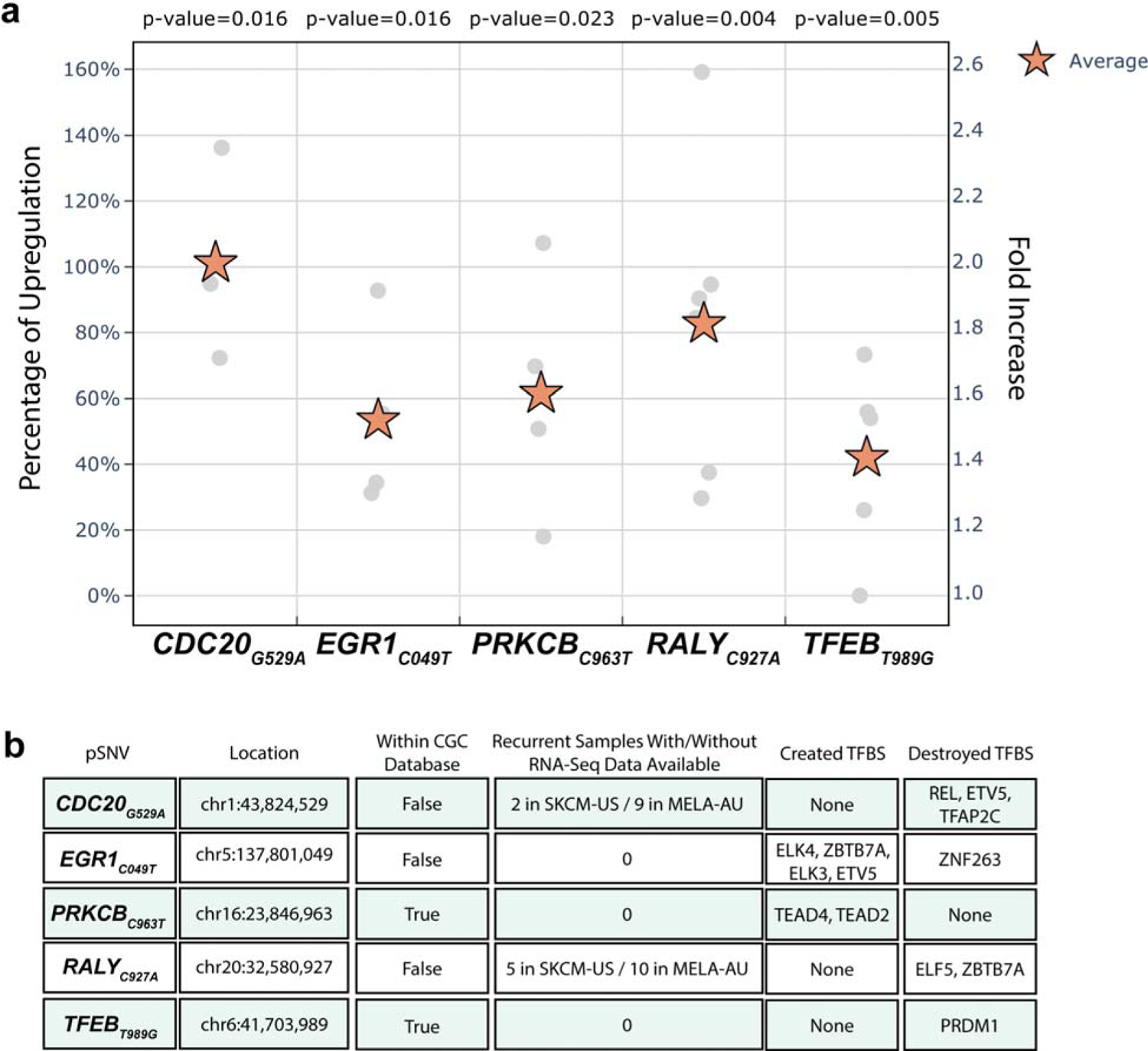
Results of the in vitro validation experiments for the 5 positively validated candidates and their key features. **(a)** A scatterplot of the percentage of upregulation and fold-increase (y-axis) for each positively validated pSNV (x-axis). An upregulation of 0% implies that the activity of the wild type is equal to the activity of the mutant (i.e. fold change of 1), whereas an upregulation of 100 % implies activity doubling (i.e. fold change of 2). The multiple replicates (gray circles), the average of these replicates (orange star), and each pSNV’s p-value using a one-sided t-test are also denoted. **(b)** A table containing the key feature s of each positively validated pSNV.

The validation rate of the recurrent (2 out of 6; 33%; *RALY_C927A_* and *CDC20_G529A_*) mutations was comparable to that of our singleton (3 out of 8; 38%; *PRKCB_C963T_*, *EGR1_C049T_* and *TFEB_T989G_*) candidates indicating a relatively equal propensity for functional significance regardless of recurrence status.

### Presence of Mutational Signatures within SKCM-US for the validated mutations

Including *ANKRD53_G529A_*, the previously-identified *CDC20* ^13,14^ and *RALY* ^2^ pSNVs, 10 of the 15 (67%) selected candidates were identified in the skin cutaneous melanoma (SKCM-US) cohort. Cutaneous melanoma, in particular, typically displays an exceptionally high mutational load^29^ that is predominantly attributed to the novel COSMIC mutational signatures SBS7a, SBS7b and SBS7c^29^ associated with ultraviolet radiation exposure.

We observed that these three mutational signatures were distinctly over-represented in comparison to all other non-melanoma cohorts in both genome-wide SNVs and pSNVs (Fig. 4; Supplementary Data 5), which agrees with previous studies. Moreover, we found that of the 10 validated SKCM-US candidates, each mutation matched to at least one of the UV-induced mutational signatures (3 matched exclusively with SBS7a, 4 match exclusively with SBS7b, and 3 matched with both SBS7a and SBS7b).

**Fig. 4:**
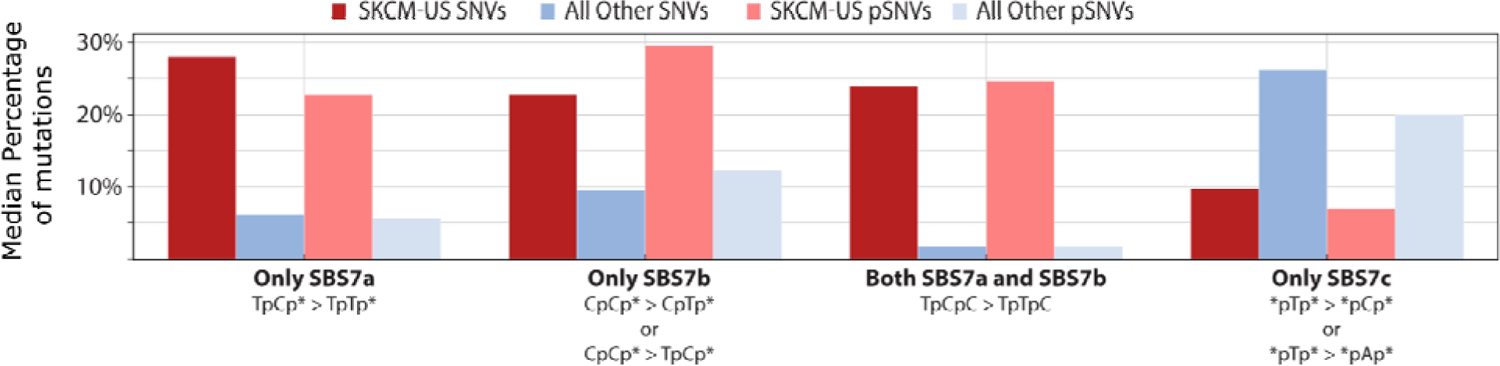
A comparison of the median percentage of mutations within all samples (y axis) belonging to the SBS7 mutational signatures (x-axis) between melanoma and non-melanoma SNVs and pSNVs. Below each mutational signature, its respective trinucleotide context (i.e. TpCp* > TpTp*) is displayed; the star (*) represents an y of the four nucleotides (i.e. TpCp* implies that this can be TpCpA, TpCpC, TpCpT, o TpCpG).

### Validating the REMIND-Cancer Approach on the EOPC-DE Cohort Prioritizes th e Proto-Oncogene *MYB*

We generated a validation dataset by integrating the RNA-seq data for 23 early onset prostate cancer (EOPC-DE) patients with their corresponding WGS data. While the WGS data was originally part of the PCAWG project, the RNA-seq data had not been previously incorporated^30^.

Of the 56,608 total EOPC-DE mutations, 554 fit our pSNV criteria (Fig. 5a). Subsequently, 54 of these mutations passed our additional filtering steps, of which only one pSNV was recurrent among the whole PCAWG dataset.

**Fig. 5:**
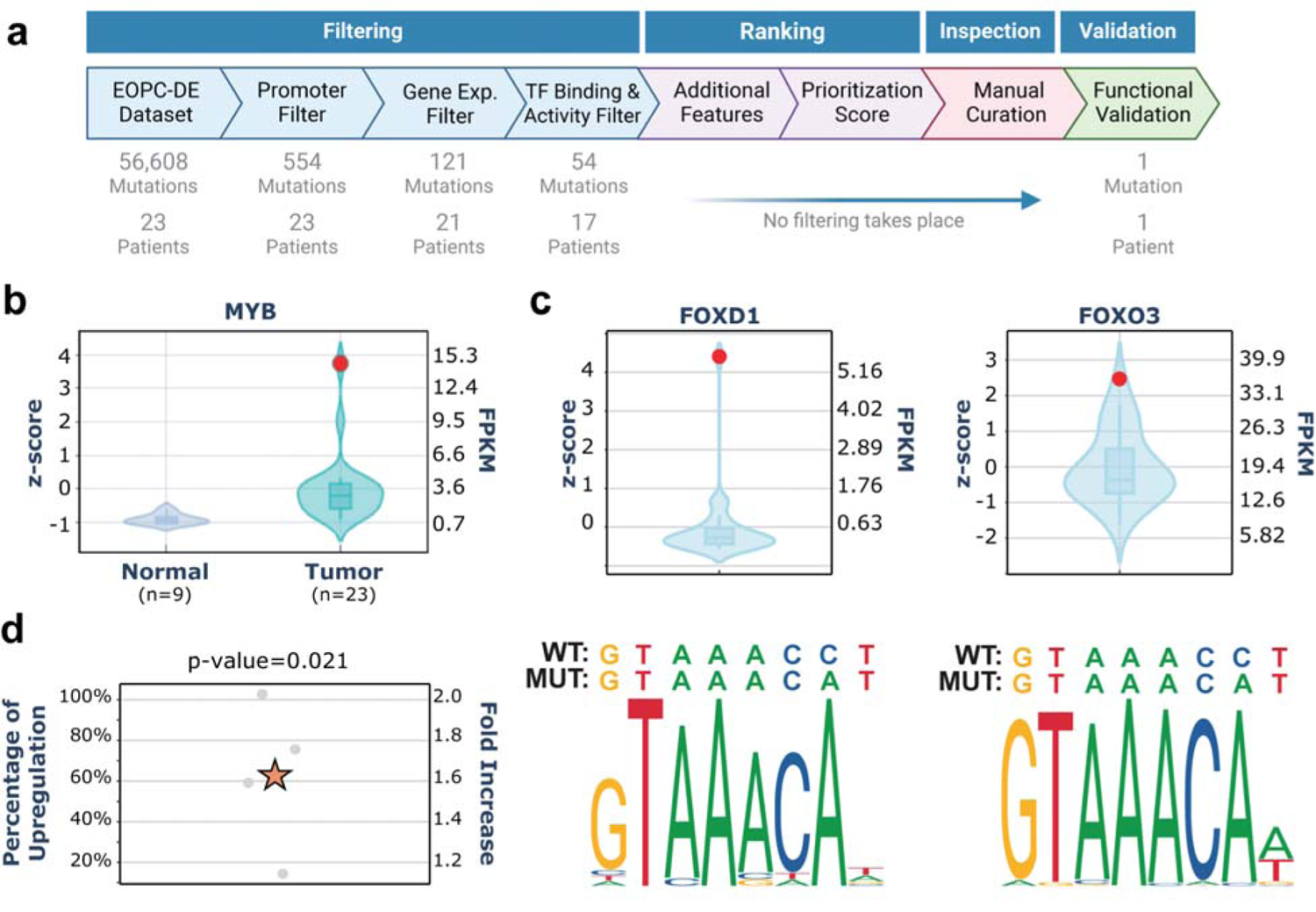
Results of applying the REMIND-Cancer workflow to the early-onset prostate cancer (EOPC-DE) cohort and of the highest-ranking pSNV in the MYB promoter. **(a)** Results during each step of the REMIND-Cancer workflow^27^ **(b)** Violin plot of the normalized (z-score; left axis) and absolute gene expression level (FPKM; right axis) for the normal and tumor samples of MYB. The red dot represents the expression in the mutated sample. **(c)** Violin plots (top) depicting the z-score (left axis) and FPKM (right axis) values of the TFs FOXD1 and FOXO 3 relative to the rest of the EOPC-DE cohort. The red dot represents the expression level of the respective TF in the patient harboring MYB _C964A_. Below the violin plots, the corresponding logo plots along with the wild type / mutant sequence for bot h TFs are depicted. **(d)** Luciferase validation results of the MYB pSNV are shown (p value = 0.021, one-sided t-test).

Here, the highest score was assigned to a singleton pSNV that created a TFBS for two forkhead box (FOX) TFs within the promoter of the proto-oncogene *MYB*. *MYB* was overall upregulated in EOPC-DE samples compared to healthy controls (Fig. 5b) and was found to be strongly upregulated in the sample harboring *MYB_C964A_* compared to the rest of the EOPC-DE cohort (z-score = 3.75). Additionally, the TFs, namely *FOXD1* and *FOXO3*, showed significantly increased expression in the original sample (Fig. 5c). In the validation experiment, *MYB_C964A_* led to a statistically significant increase in luciferase activity of 63% (Fig. 5d; one-sided t-test; p-value = 0.021).

## Discussion

In this study, we introduce the *Regulatory Mutation Identification ‘N’ Descriptions in CANCER* (REMIND-Cancer) workflow, which is an integrative approach to characterize and identify functional promoter pSNVs that lead to an upregulation of their associated genes. It follows a filter-ranking-inspection-validation paradigm, in which a series of filtering steps are applied to pSNVs. Biologically-relevant features are considered for the remaining mutations, and these genomic, transcriptomic and annotation-based features are used to compute a prioritization score for subsequent ranking. *DeepPileup* provides information on the distribution of potential technical artifacts at the mutated positions. Available copy number variation calls are used to analyze convergent cancer evolution patterns through *GenomeTornadoPlot*, which is especially useful for genes not yet functionally implied in cancer. To assist in candidate pSNV selection for *in vitro* validation, we developed *pSNV-Hunter*, a visualization tool for inspection of the high-scoring candidates. Moreover, we have established an experimental workflow such that the functionality of these candidate mutations can be tested using a luciferase reporter assay. By comparing activity levels between the wildtype and mutant, we can discern the effect of specific candidate mutations, further enhancing our understanding of their functional implications.

With the computed prioritization score as well as through our manual curation efforts, we identified pSNVs within *ANKRD53, MYB* and 14 additional candidate promoter mutations for further *in vitro* validation. While *ANKRD53* has not yet been implicated directly in cancer, depletion of this protein delays progression of mitosis and interferes with chromosome alignment^31^, which potentially ties its expression level to the proliferation rate of cancer cells. Notably, the introduction of *ANKRD53_G529A_* did not cause a statistically significant upregulation of luciferase reporter activity on its own. Yet, stimulation with TNF-α to activate *NF-kB*, the TF predicted to bind to the newly-created TFBS, resulted in a significant increase in promoter activity. This emphasizes the relevance of the model system used. While we carefully ensured that the TFs predicted to bind to the altered TFBS were expressed in HEK293FT cells based on RNAseq data from the Human Protein Atlas, it was out of scope to validate the activation state of all selected candidate TFs. Beyond *ANKRD53_G529A_*, 6 (*MYB_C964A_*, *CDC20_G529A_*, *RALY_C927A_*, *PRKCB_C963T_*, *EGR1_C049T_*, and *SECISBP2_G357A_*) of the 15 selected pSNVs showed a statistically significant upregulation (p-value ≤ 0.05) in the HEK293FT kidney cell line in the absence of pathway stimulation. Together, these findings highlight the potential of our computational pipeline to identify functional pSNVs. Moreover, they underscore the functional impact of the identified pSNVs and their potential relevance.

With a FPKM z-score of 3.75, the singleton *MYB_C964A_* pSNV was associated with strong transcriptional upregulation in the respective prostate cancer patient and also showed significant activation in the luciferase assay. While this proto-oncogene has mainly been associated with differentiation of hematopoietic cells and its role in leukemia, there are indications of a role in prostate cancer as well^32,33^. Specifically, amplification and the resulting overexpression of *MYB* was associated with the acquisition of a hormone-resistant phenotype^34^. Furthermore, ectopic overexpression of *MYB* in LNCaP cells caused androgen-deprivation resistance and induced PSA (prostate specific antigen) expression^35,36^. Potentially, this phenotype can be attributed to a direct interaction between *MYB* and the androgen receptor (AR) to sustain AR localization inside the nucleus^35,37^. Together, these studies point at a potential selective advantage of prostate cancer cells induced by overexpression of *MYB*. In the light of these findings, *MYB_C964A_* might display an alternative genetic mechanism towards the upregulation of *MYB* in the absence of genomic amplification, indicating a case of convergent tumor evolution. Moreover, this case exemplifies how the potential genetic cause for resistance to an established therapy may be overlooked due to being a singleton mutation.

When considering all validated candidate mutations (*ANKRD53_G529A_* + 14 additional candidates + *MYB_C964A_*), the statistically significant recurrent mutations (namely *ANKRD53_G529A_, CDC20_G529A_*, and *RALY_C927T_*) positively validated at a rate of 3/7 (43%) comparable to that of the statistically-significant singletons, namely *PRKCB_C963T_*, *EGR1_C049T_*, *TFEB_T989G_*, and *MYB_C964A_* (4/9; 44%). This observation provides compelling evidence that less frequent functional promoter mutations that appear as singletons in small and mid-sized cohorts can be identified using the REMIND-Cancer pipeline.

Therefore, the workflow is part of the answer to the demand for the characterization of rare driver mutations that has been raised within the precision oncology field^38^. The similar rates of positive validation in both recurrent and singleton mutations underscore the importance of examining the functional impact of a diverse range of mutations whether recurrent or not. Rare cancer types constitute about 25% of all cancer cases and are responsible for a quarter of all cancer related deaths^39^. For these cancer types, recurrence-independent approaches, such as the REMIND-Cancer workflow, are required as these cohorts are notoriously statistically underpowered.

Previously, two studies have claimed adverse effects of the *CDC20_G529A_* pSNV. One study claims that an *increase* in the transcription of this gene was achieved through the disruption of the binding of the repressor ELK4^13^. By luciferase reporter assay, a significant upregulation was reported in the kidney cell line HEK293 while a consistent but insignificant effect was observed in the melanoma cell line M14. However, in another study, a statistically-significant *reduction* in luciferase activity within HEK293 was reported^14^. Within our study, we also functionally validated the recurrent *CDC20*_G529A_ pSNV within the HEK293 cell line and observed a significant upregulation, agreeing with He et al.^13^. More specifically, our results showed a more pronounced luciferase activity compared to the previously-reported 50%.

We posit that the difference in these results are attributable to the length and positioning of the synthesized promoter sequence used in the luciferase construct (He *et al.* 859 bp construct^13^; Godoy *et al*. 170 bp construct^14^; REMIND-Cancer 663 bp construct), which influences the distance to the TSS and the presence of co-regulatory sites. However, to conclusively demonstrate the impact of a pSNV, it needs to be recapitulated within its native chromatin context e.g. by CRISPR-based technologies to investigate target gene upregulation and transcription factor binding in an isogenic context.

Among the 16 pSNVs that were selected for *in vitro* validation, 10 were derived from the skin cutaneous melanoma cohort (SKCM-US). Of those mutations, only *RALY_C927T_* has previously been identified as being a potential driver mutation^29^ but this particular mutation has not been validated until now.

To our knowledge, other than *CDC20_G529A_* and *RALY_C927T_*, no other candidate pSNVs described here have been previously characterized. Hence, our study advances the repertoire of functionally validated pSNVs with a potential role in cancer.

In this study, we addressed the question if the lack of established activating promoter mutations beyond *TERT* in cancer are due to mutation bias, selection bias or method bias. In other words, is the creation of promoter SNVs just less frequent and therefore not captured by current methods or is it not occurring due to a lack of frequently mutated targets providing a substantial selective advantage? In this context, target refers to a nucleotide in a promoter that, by a single point mutation, leads to an upregulation of the adjacent gene.

Our observation in SKCM-US indicates that mutation bias indeed is a part of the explanation as the UV-related signature SBS7a and SBS7b generate an abundance of cytosine mutations. Additionally, unlike the ubiquitous age-associated spontaneous deamination SBS1, SBS7 affects cytosines outside a CpG context. The abundance and specific mechanism of mutations in melanoma results in mutations that are otherwise less frequent in cancer. The consequences of this process is also reflected in the frequency of the two already established *TERT* promoter mutations that can also be attributed to SBS7a and display, among all cancer types, the highest frequency in cutaneous melanoma (82%)^40^.

Additionally, *ANKRD53_G529A_* and *MYB_C964A_* indicate a selection bias. In the case of *ANKRD53_G529A_,* the pSNV was necessary but not sufficient for its upregulation due to the requirement of an activation of the RELA-encoded transcription factor p65. In the case of *MYB_C964A_,* two TFs, particularly FOXD1 and FOXO3, which were predicted to bind the *de novo* motif, displayed a strong upregulation in the original patient sample compared to its cohort. Both of these mutations only become effective in cells or tissues that either naturally provide the necessary conditions or rely on a second hit that establishes these conditions during the evolution of the cancer. This conditional selective pressure can be interpreted as selection bias.

Lastly, it is uncertain if the statistical power to identify lowly recurrent non-coding driver mutations may ever be sufficient in datasets with limited sample size. Our analysis shows that the REMIND-Cancer workflow can identify novel functional pSNVs in the PCAWG dataset, which has been previously analyzed intensively by many state-of-the-art methods for non-coding driver identification^8^. This indicates a blind spot for lowly and non-recurrent regulatory functional mutations and thus a method bias.

Although it is doubtful that other events on-par with the pan-cancer relevance of the *TERT* pSNVs can be found among this long tail of less frequent pSNVs, some of these cases may constitute the hidden co-driver events that explain variable response to targeted therapies and the yet undiscovered causes of some rare cancers. To move from analyzing small subgroups towards achieving truly personalized oncology in the future, precision oncology programs must assess singletons in order to decrypt the dark matter of the cancer genome.

## Online Methods

The REMIND-Cancer workflow follows a filtering-ranking-inspection-validation paradigm in which patients and their corresponding somatic pSNVs are (1) filtered to attempt to focus only on putative driver mutations, (2) scored and ranked based on genomic, transcriptomic, and annotation features, and (3) manually inspected to remove artifacts and ensure relevance for (4) validation by luciferase assay.

### Promoter Filter

Only mutations within promoter regions were analyzed. To be more inclusive than previous studies^8^, the promoter region was defined as ranging from 1,000 base pairs upstream to 500 base pairs downstream of the transcription start site (TSS). To be consistent with the PCAWG project, all coordinates are based on the GRCh37 (hg19) genome assembly and the GENCODE annotation V19^41^ and only the canonical transcripts were considered. A list of the used promoter definitions can be found in Supplementary Data 6.

### Gene Expression Filter

To identify upregulated genes, we utilized the RNA-Seq-based gene expression data available for 927 samples and subsequently disregarded pSNVs from samples lacking RNA-Seq data. The gene expression, measured in fragments per kilobase of transcript per million mapped reads (FPKM), of the gene adjacent to the mutated promoter was z-score normalized relative to samples within its subcohort. A z-score greater than 0 then indicated an above-average expression and the corresponding pSNVs were retained.

### TFBS Motif and Expression Filter

Mutations that impacted the binding motif of transcription factors were identified within this filter. Utilizing the Find Individual Motif Occurrences (FIMO) tool^42^ from the MEME Suite toolkit^43^ along with the JASPAR2020^44^ database of curated transcription factors, the DNA sequence of +/- 10 bp around every pSNV underwent scanning by FIMO against every TF motif in JASPAR2020. Using the positional weight matrix of the TF and its associated sequence context, FIMO calculated a statistical binding affinity score, which indicates the likelihood of observing the specific motif. To assess the impact of a mutation, the ratio (denoted as S(TFBS)) between the binding affinity scores for the mutant over the wild type allele was calculated. This ratio serves as a quantitative measure indicating the impact of mutations on the binding motif of TFs. To determine this effect, we established a threshold criteria of S(TFBS)>11 for a created TFBS motif and S(TFBS)<0.09 for a destroyed TFBS motif. Mutations were kept if at least one predicted TF fulfilled one of these criteria.

Next, we used the RNA-seq data of the patients harboring the pSNV to obtain the expression of the predicted TF. pSNVs were retained if it altered at least one TFBS for which the matching TF had a FPKM > 0.

### Recurrence, CGC, and Chromatin Accessibility

Additional pSNV features were annotated for the computation of the prioritization score and subsequent ranking. Of these features, recurrence was determined by utilizing all available WGS data irrespective of RNA-Seq availability, which encompassed 19,401,901 mutations from 2,413 samples. Specifically, recurrence was determined by creating an index of all pSNVs based on their chromosome, position and nucleotide change using a Python dictionary. The number of entries for each locus was queried to obtain a recurrence statistic. Furthermore, pSNVs matched to genes listed as one of the 746 entries in the Cancer Gene Census (CGC) database^45^ as of 11 November 2020 were labeled as CGC “True”. All CGC tiers of evidence were equally considered for the prioritization scoring (Supplementary Data 6). To annotate the presence of open chromatin overlapping with the pSNV, their positions were intersected with the ChromHMM annotation as found in the ‘hg19_genome_100_segments’ file^46^ which was last modified on 5 December 2022 (Supplementary Data 7). These chromatin-state annotations were based on their universal full-stack model, which was created by training a hidden markov based model on 1,000+ datasets within 100+ cell types. Any pSNV labeled as being within the regions of a bivalent promoter (i.e. BivProm1-4) or containing one of the histone modifications H3K4me1-3 (i.e. PromF1-5) were labeled as having open chromatin. The

### Candidate Mutation Prioritization

To prioritize mutations for *in vitro* functional validation, an empirically calibrated weighted-sum scoring function was used to compute a prioritization score. The function favorably scored pSNVs with strong upregulation, in regions of open chromatin, from samples with high tumor purity and with high allele frequency indicating clonality. Further, recurrence in other samples and association to cancer genes increased the score and, to a degree, the mutation of multiple TFBSs was acknowledged.

The specific genomic, transcriptomic, and annotation features that were used and their weights can be found in Supplementary Fig. 1. We calibrated the weights to prioritize specificity over sensitivity. For recurrence and the number of created/destroyed TFBSs, a maximum contribution value to the score is given. Additionally, for tumor purity and allele frequency, a boolean feature weight was added to the prioritization score if they met or exceeded their specific threshold value. Lastly, when scoring both open chromatin at a pSNV position and presence of the adjacent gene within the CGC list, a fixed weight was added to the score if the features were “True”. More specifically, the prioritization score was computed using the following formula:

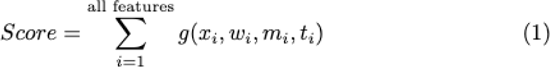

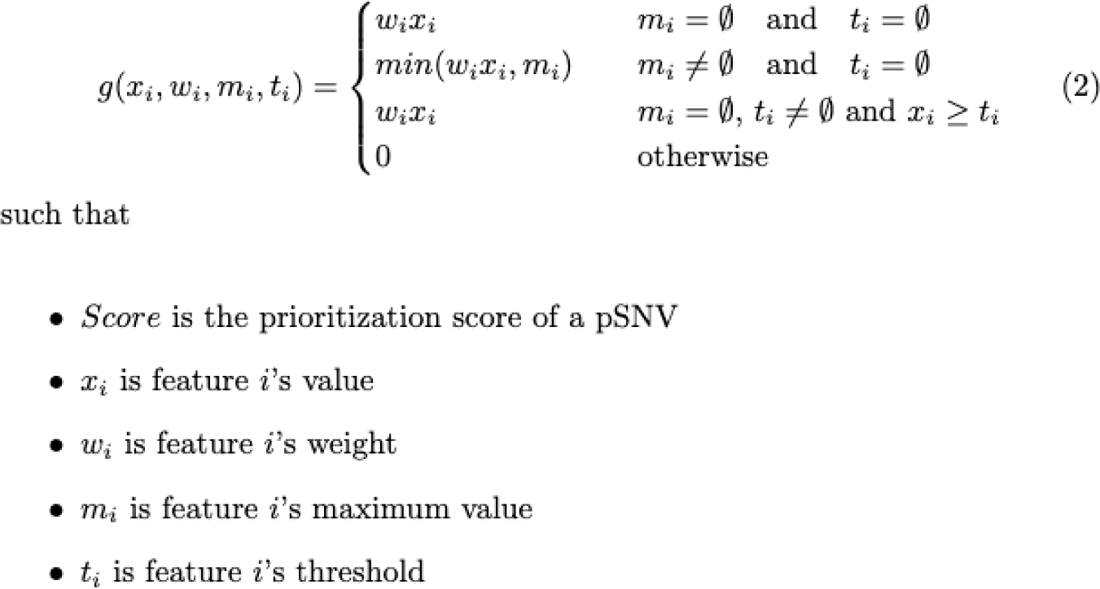

### Inspection Step

As quality control measures, we implemented two established tools, *DeepPileup*^8^ and *GenomeTornadoPlot*^47^ prior to *in vitro* validation. *DeepPileup* tests if the genomic positions of a SNV is prone to report sporadic variant repeats by analyzing pileups in the tumor and control samples of the full PCAWG dataset. Then, the percentage of samples per cohort in which more than two variants (2V+) are observed is visualized. Germline variants are identified as positions with minor allele frequency > 25% in control samples. Remaining candidates in which the control samples reached a similar or higher 2V+ percentage than tumor samples in more than two cohorts were considered noisy and not selected for validation. An example of a noisy genomic position can be seen in Supplementary Fig. 2.

Further, for each pSNV-gene pair we employed the *GenomeTornadoPlot*^47^ package to test for an enrichment of focal high level amplifications in the remaining PCAWG cohort indicative of an alternative gene activation mechanism in other tumors.

### pSNV Hunter

To assist in the inspection step when selecting pSNVs for *in vitro* validation, we created pSNV Hunter, a multi-functional software tool that allows for the comprehensive examination of individual pSNVs. Through an interactive dashboard that was created in Python using the Plotly Dash package, users can view plots displaying the gene expression of the adjacent gene corresponding to the pSNV, recurrence level, and TF expression while also viewing specific gene and TF function provided by the NCBI. Both sample-specific information and quality control plots from *DeepPileup* and *GenomeTornadoPlot* can also be seen. At the time of this publication, pSNV Hunter is in its beta stage but has been made publicly-available on GitHub at: https://github.com/nicholas-abad/pSNV-hunter.

### Software Used

The REMIND-Cancer pipeline was run using Python 3.10.11 and primarily utilized the following packages: Pandas (2.0.1)^48^, NumPy (1.24.3) ^49^ and SciPy (1.11.2)^50^. The full list of Python packages can be found within our publicly-available GitHub Repository, which can be found in *Online Methods: Code Availability*. Two stand-alone wrappers for *DeepPileup* (Python 3.10.11) and *GenomeTornadoPlot* (R 4.0.0) were created to output results in a format that was compatible with the REMIND-Cancer Pipeline. The *DeepPileup* wrapper can be found at https://github.com/nicholas-abad/deep-pileup-wrapper whereas the *GenomeTornadoPlot* wrapper can be found at https://github.com/nicholas-abad/genome-tornado-plot-wrapper.

### Luciferase reporter assays

Wildtype and mutant promoter fragments were inserted into the pCR8_GW_luc plasmid (kindly provided by Dr. Rainer Will) via gene synthesis (BioCat, Heidelberg, Germany). Sequences and restriction enzymes used for subcloning are listed in Supplementary Data 8. HEK293FT cells (ATCC PTA-5077) were maintained in DMEM medium (Gibco, #41966-29) supplemented with 10% FBS (Gibco), 1% L-Glu (PAN BIOTECH, #P04-80100), 1% NEAA (Gibco, #11140-039) and 500µg/ml Geneticin (Gibco, #10131-027) at 37°C and 5% CO_2_ and subcultured twice a week at a confluence of around 70-80%. Cells were tested for potential mycoplasma contamination on a regular basis, and were positively authenticated prior to and at the end of the study (Multiplexion GmbH, Heidelberg, Germany).

For luciferase reporter assays, cells were seeded in 96-well plates (Greiner bio-one, cellstar #655180) at 7,500 cells/well in growth media and 24h later transfected with 90 ng/well pCR8 and 10 ng/well pRL-CMV (Promega, #E2261) using 0.4 µl/well Lipofectamine-2000 (Thermofisher) in 100 µl/well OptiMEM (Gibco, #31985-047). The medium was changed 5h post transfection to full media and cells were incubated for 48h before luciferase activities were measured as described previously^51^ using a GloMax Discover microplate reader (Promega GmbH, Walldorf, Germany). For siRNA knockdown, 3nM of a complex (30-plex) siRNA from siTOOLs (Plenegg, Germany) were co-transfected together with the plasmid mix. In case of stimulation with TNF-α (PeproTech, # 300-01A) cells were incubated with 20 ng/ml for 5h prior to lysis and luciferase assay.

For each independent biological repeat, multiple technical replicates (i.e. multiple identically transfected wells) were prepared per condition. Lysates were transferred to white 96-well plates (Greiner bio-one, #655083). For each well, the signal obtained from firefly luciferase was normalized to the Renilla luciferase signal and the median of the six technical replicates was recorded. The relative change in activity of the mutant promoter was calculated as the change in normalized signal between the mutant and the corresponding wildtype vectors in percent and plotted for each independent biological replicate (n≥3).

### Mutational Signature Detection

The presence of each of the three UV-induced mutational signatures (SBS7a, SBS7b, and SBS7c^52^) were annotated for the remaining mutations belonging to the SKCM-US cohort using only the base context.

In the case of SBS7a, we determined that a pSNV matched this signature if the pSNV was a C>T mutation in a TpC dinucleotide context (i.e. TC > TT), representing 75% of the SBS7a cases in the COSMIC database. For SBS7b, a pSNV contained this signature if there was a C>T mutation at a CpC dinucleotide (i.e. CC>CT or CC>TC; 71% of all COSMIC SBS7b cases) and lastly, a pSNV contained SBS7c if it was either a T>C or T>A mutation (72% of all COSMIC SBS7c cases).

### PCAWG Dataset

Both the WGS and RNA-Seq data used in this study were from an internal 2015 PCAWG data freeze. To match a sample’s WGS to their RNA-Seq data, the specimen’s submission ID (WGS) was matched to its corresponding submitter bundle ID (RNA-Seq). Most WGS cancer types had corresponding RNA-Seq data available (Supplementary Fig. 5).

### EOPC-DE expression data

The ICGC’s early onset prostate cancer (EOPC-DE) WGS data from PCAWG was integrated with the gene expression data^30^.

### Statistical Testing

Statistical tests of the luciferase assay results comparing the percentage of upregulation between the wild type and mutant were conducted using a one-sided t-test with an expected value of 0. Explicitly, the null hypothesis (H_0_) is that the mean percentage upregulated in the mutant is less than or equal to the mean percentage upregulated in the wild type. Therefore, a significant p-value (i.e. p-value ≤ 0.05) implies rejecting the null hypothesis, providing evidence of a positive upregulation in the mutant compared to the wild type.

## Data Availability

PCAWG data can be downloaded in the PCAWG data portal (https://dcc.icgc.org/pcawg).

## Code Availability

The publicly-available REMIND-Cancer pipeline code can be downloaded at https://github.com/nicholas-abad/REMIND-Cancer.

## Competing Interests

There are no competing interests.

## Acknowledgements

We thank Dr. Rainer Will, Melanie Weis and Marion Bähr for their helpful comments towards the project as well as Sabine Karolus for excellent technical support. We also thank Prof. Dr. Stefan Wiemann for his comments regarding the manuscript. NA has been supported by the Fritz Thyssen Stiftung’s REMIND Cancer grant and LF has been supported by the German Federal Ministry of Research and Education through the EOPC-Data Mining Grant (01KU1505A).

## Supplement

**Supplementary Fig. 1:**
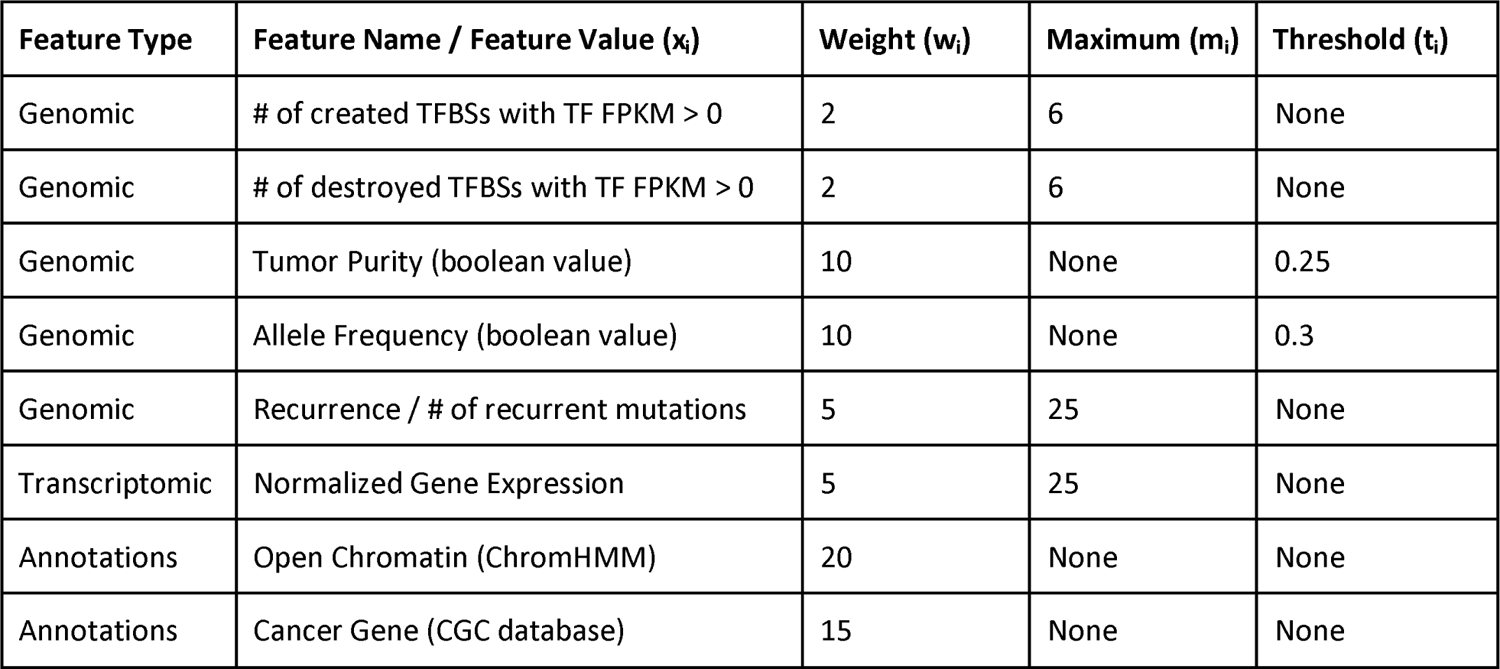
A list of the features and their corresponding weights that are considered when computing the weighted-sum prioritization score. Specifically, ‘Feature Typè refers to one of the three (genomic, transcriptomic, and annotations) types of features considered, ‘Feature Namè refers to the name/description of the feature, ‘Weight’ refers to the weight given during the computation, ‘Maximum’ refers to the maximum contribution (i.e. max(maximum value noted, x_i_ * w_i_)) that the feature could contribute to the prioritization score (if applicable), and ‘Threshold’ refers to the threshold needed to contribute to the prioritization score (if applicable).

**Supplementary Fig. 2:**
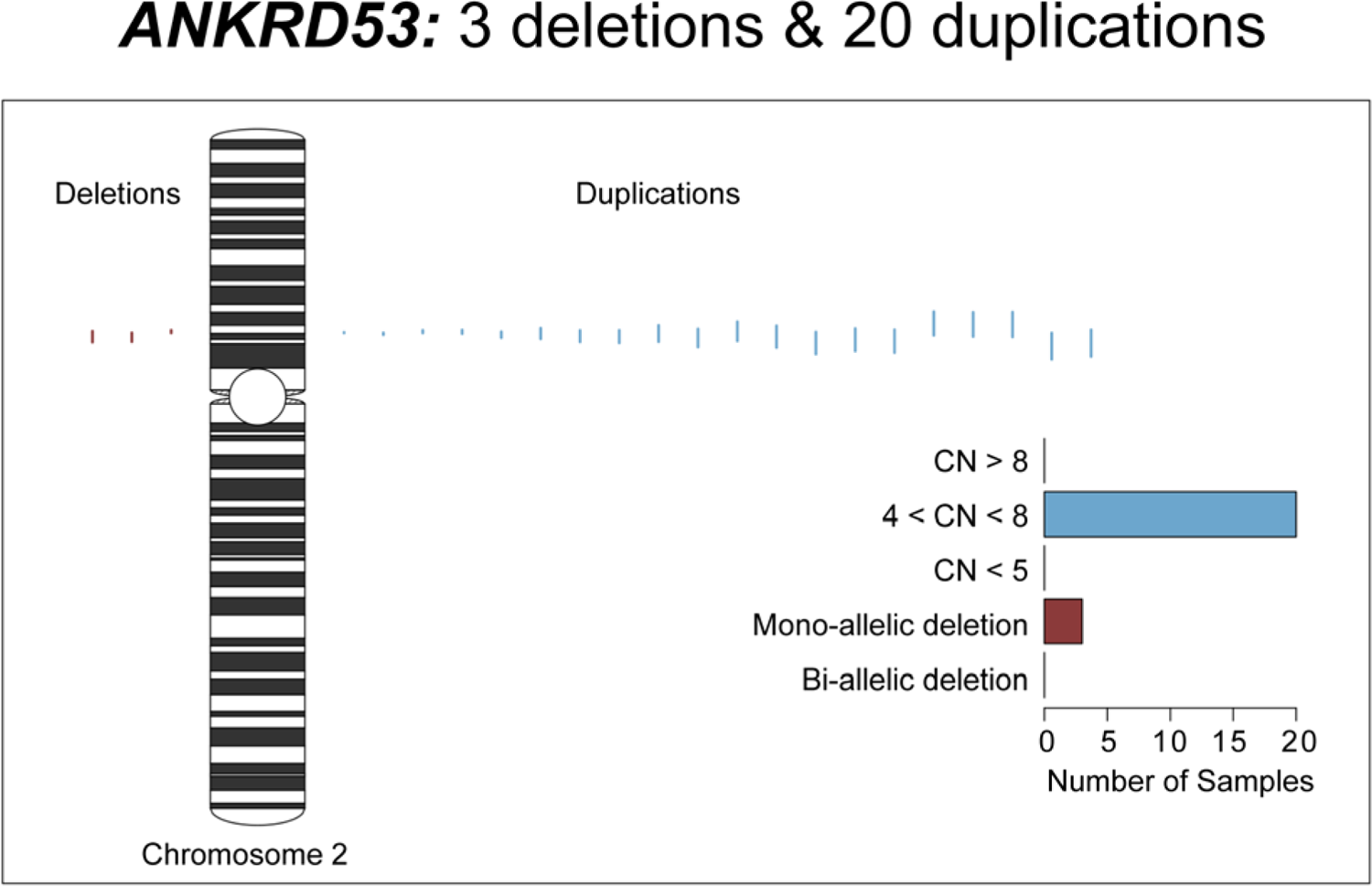
The GenomeTornadoPlot for the highest-ranking ANKRD53 pSNV within the entire PCAWG cohort. All focal deletions (3) and duplications (20) of the PCAWG dataset that overlap the gene are displayed along a schematic of the chromosome. Here, focal CNVs are defined to be shorter than 10 Mbp. Furthermore, the bar plot shows the specific copy number levels of these patients.

**Supplementary Fig. 3:**
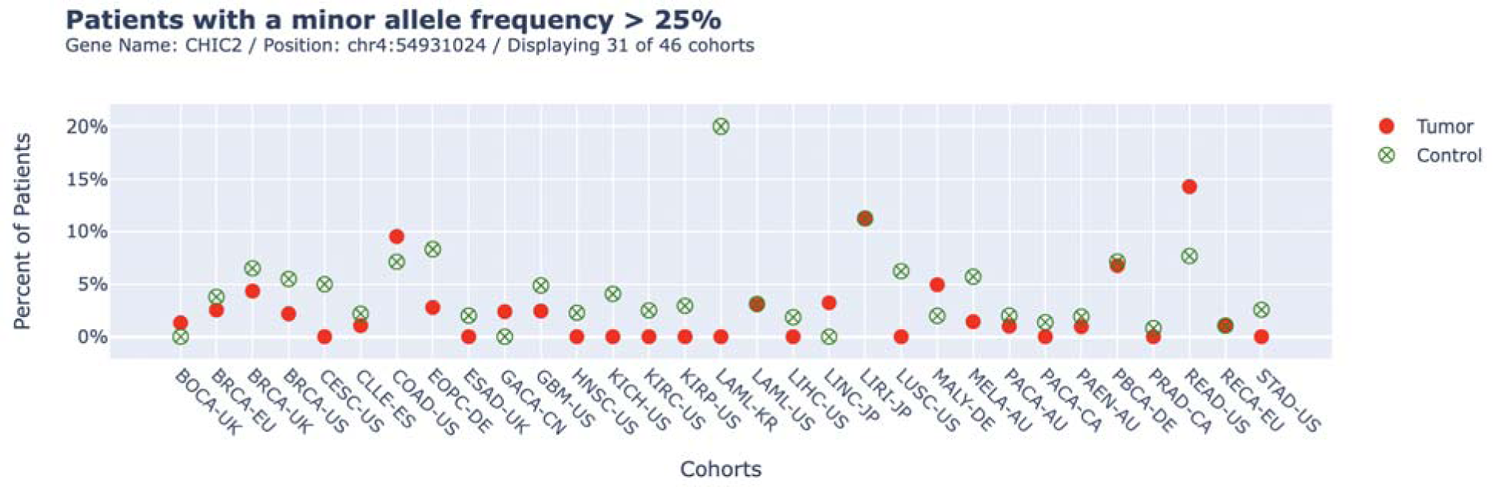
An example of a diagnostic plot generated by DeepPileup for a noisy genome position. Visualizing the CHIC2 pSNV (chr4:54931024 A>G; Rank 277), the diagram raised questions about its authenticity as it appeared t o be a potential artifact. Across various cohorts’ control samples, such as ESAD-UK, HNSC-US, KIRC-US, a substantial number of samples have more than 25% of al l reads at the locus reporting a variant. In several additional cohorts, both control and tumor samples exhibited a signal, including BRCA-EU, BRCA-UK, COAD-US, among others. Consequently, this pSNV was deemed unreliable.

**Supplementary Fig. 4:**
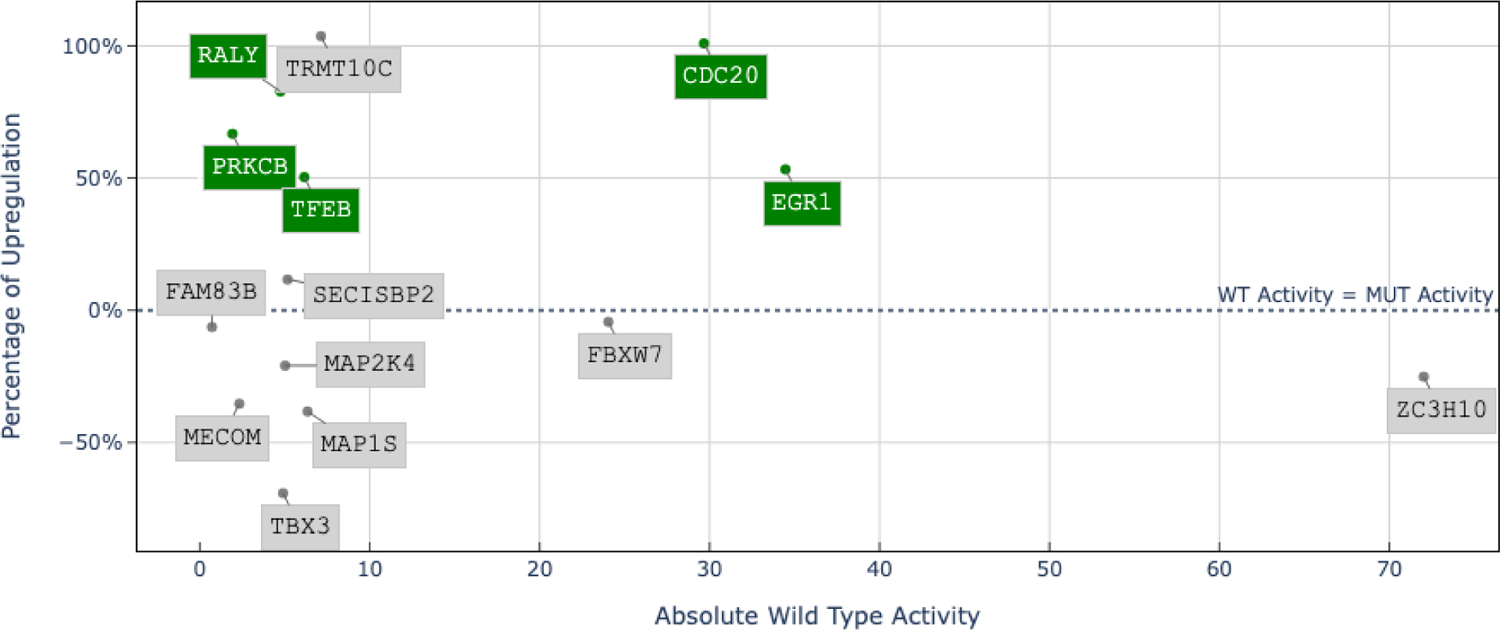
Luciferase reporter activity results for the 14 validate d pSNVs. The percentage of upregulation is on the y-axis, whereas the wild type activity is on the x-axis. The 5 positively validated pSNVs are in green.

**Supplementary Fig. 5:**
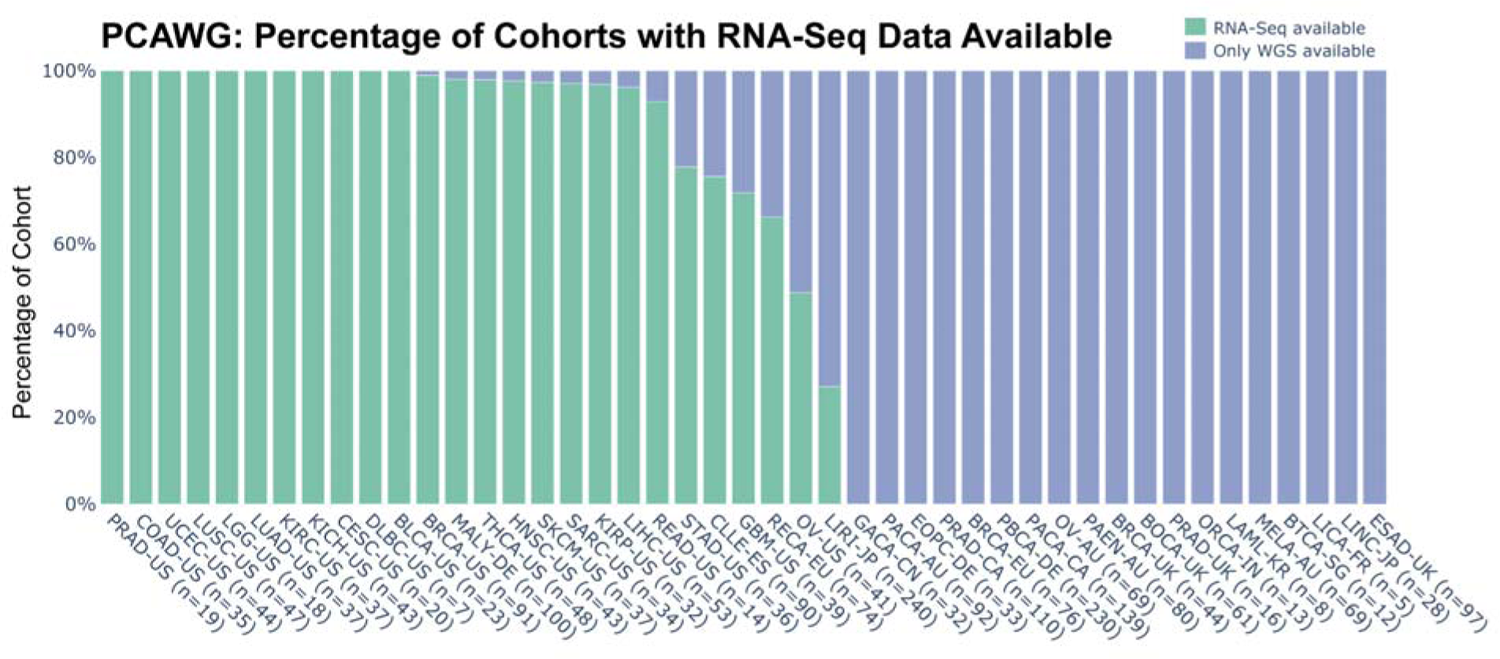
A bar plot representing the percentage of patients in each cohort that also have RNA-Seq data available within the PCAWG dataset. A value of 100% (i.e. PRAD-US, KIRC-US, etc.) implies that every WGS sample has a corresponding RNA-Seq file. A majority of cohorts (i.e. CMDI-UK, PRAD-UK, etc.) did not have any samples with corresponding RNA-Seq data available and therefore did not pass the entirety of the REMIND-Cancer pipeline.

## Supplementary Data to Provide

⍰ Supplementary Data 1: Top 1,000 ranking pSNVs (.vcf file)
⍰ Supplementary Data 2: All promoter fragments
⍰ Supplementary Data 3: ANKRD53 Luciferase Results
⍰ Supplementary Data 4: Validated Mutations Data

⍰ 4A: Luciferase Assay Percentage Upregulation
⍰ 4B: Raw Luciferase Assay Results
⍰ 4C: Validated results (.vcf file)
⍰ Supplementary Data 5: Table of SKCM statistics
⍰ Supplementary Data 6: Genomic locations of promoters used with TSS
⍰ Supplementary Data 7: ChromHMM / Regions with Open Chromatin
⍰ Supplementary Data 8: Sequence and restriction enzymes
⍰ Supplementary Data 9: Mutations per filter step

## Notes

### Competing Interest Statement

The authors have declared no competing interest.

## References

1. Suehnholz, S. P. et al. Quantifying the Expanding Landscape of Clinical Actionability for Patients with Cancer. Cancer Discov. 14, 49–65 (2024).

2. O’Dwyer, P. J. et al. The NCI-MATCH trial: lessons for precision oncology. Nat. Med. 29, 1349–1357 (2023).

3. Min, H.-Y. & Lee, H.-Y. Molecular targeted therapy for anticancer treatment. Exp. Mol. Med. 54, 1670–1694 (2022).

4. Zhou, Z. & Li, M. Targeted therapies for cancer. BMC Med. 20, 90 (2022).

5. Zhong, L. et al. Small molecules in targeted cancer therapy: advances, challenges, and future perspectives. Signal Transduct. Target. Ther. 6, 1–48 (2021).

6. Aaltonen, L. A. et al. Pan-cancer analysis of whole genomes. Nature 578, 82–93 (2020).

7. Martínez-Jiménez, F. et al. A compendium of mutational cancer driver genes. Nat. Rev. Cancer 20, 555–572 (2020).

8. Rheinbay, E. et al. Analyses of non-coding somatic drivers in 2,658 cancer whole genomes. Nature 578, 102–111 (2020).

9. Horn, S. et al. TERT Promoter Mutations in Familial and Sporadic Melanoma. Science 339, 959–961 (2013).

10. Huang, F. W. et al. Highly Recurrent TERT Promoter Mutations in Human Melanoma. Science 339, 957–959 (2013).

11. Li, C. et al. The C228T mutation of TERT promoter frequently occurs in bladder cancer stem cells and contributes to tumorigenesis of bladder cancer. Oncotarget 6, 19542–19551 (2015).

12. Bullock, M. et al. TERT promoter mutations are a major indicator of recurrence and death due to papillary thyroid carcinomas. Clin. Endocrinol. (Oxf.) 85, 283–290 (2016).

13. He, Z. et al. Pan-cancer noncoding genomic analysis identifies functional *CDC20* promoter mutation hotspots. iScience 24, 102285 (2021).

14. Godoy, P. M. et al. Functional analysis of recurrent CDC20 promoter variants in human melanoma. *Commun*. Biol. 6, 1–17 (2023).

15. Fredriksson, N. J., Ny, L., Nilsson, J. A. & Larsson, E. Systematic analysis of noncoding somatic mutations and gene expression alterations across 14 tumor types. Nat. Genet. 46, 1258–1263 (2014).

16. Weinhold, N., Jacobsen, A., Schultz, N., Sander, C. & Lee, W. Genome-wide analysis of noncoding regulatory mutations in cancer. Nat. Genet. 46, 1160–1165 (2014).

17. Rheinbay, E. et al. Recurrent and functional regulatory mutations in breast cancer. Nature 547, 55–60 (2017).

18. Kim, K. et al. Chromatin structure–based prediction of recurrent noncoding mutations in cancer. Nat. Genet. 48, 1321–1326 (2016).

19. Fu, Y. et al. FunSeq2: a framework for prioritizing noncoding regulatory variants in cancer. Genome Biol. 15, 480 (2014).

20. Lochovsky, L., Zhang, J., Fu, Y., Khurana, E. & Gerstein, M. LARVA: an integrative framework for large-scale analysis of recurrent variants in noncoding annotations. Nucleic Acids Res. 43, 8123–8134 (2015).

21. Bailey, M. H. et al. Comprehensive Characterization of Cancer Driver Genes and Mutations. Cell 173, 371–385.e18 (2018).

22. Dietlein, F. et al. Identification of cancer driver genes based on nucleotide context. Nat. Genet. 52, 208–218 (2020).

23. Mularoni, L., Sabarinathan, R., Deu-Pons, J., Gonzalez-Perez, A. & López-Bigas, N. OncodriveFML: a general framework to identify coding and non-coding regions with cancer driver mutations. Genome Biol. 17, 128 (2016).

24. Gonzalez-Perez, A. & Lopez-Bigas, N. Functional impact bias reveals cancer drivers. Nucleic Acids Res. 40, e169 (2012).

25. Zhou, J. & Troyanskaya, O. G. Predicting effects of noncoding variants with deep learning–based sequence model. Nat. Methods 12, 931–934 (2015).

26. Zhou, J. et al. Deep learning sequence-based ab initio prediction of variant effects on expression and disease risk. Nat. Genet. 50, 1171–1179 (2018).

27. Adapted from ‘Icon Pack - Timelines (Horizontal)’, by BioRender.com (2024). Retrieved from Adapted from “FullTemplateName”, by http://BioRender.com (CurrentYear). Retrieved from https://app.biorender.com/biorender-templates.

28. Karolchik, D. et al. The UCSC Genome Browser Database. Nucleic Acids Res. 31, 51– 54 (2003).

29. Hayward, N. K. et al. Whole-genome landscapes of major melanoma subtypes. Nature 545, 175–180 (2017).

30. Gerhauser, C. et al. Molecular Evolution of Early-Onset Prostate Cancer Identifies Molecular Risk Markers and Clinical Trajectories. Cancer Cell 34, 996–1011.e8 (2018).

31. Kim, S. & Jang, C.-Y. ANKRD53 interacts with DDA3 and regulates chromosome integrity during mitosis. Biochem. Biophys. Res. Commun. 470, 484–491 (2016).

32. Pattabiraman, D. R. & Gonda, T. J. Role and potential for therapeutic targeting of MYB in leukemia. Leukemia 27, 269–277 (2013).

33. Nakata, Y. et al. c-Myb contributes to G2/M cell cycle transition in human hematopoietic cells by direct regulation of cyclin B1 expression. Mol. Cell. Biol. 27, 2048–2058 (2007).

34. Edwards, J., Krishna, N. S., Witton, C. J. & Bartlett, J. M. S. Gene amplifications associated with the development of hormone-resistant prostate cancer. Clin. Cancer Res. Off. J. Am. Assoc. Cancer Res. 9, 5271–5281 (2003).

35. Srivastava, S. K. et al. MYB interacts with androgen receptor, sustains its ligand-independent activation and promotes castration resistance in prostate cancer. Br. J. Cancer 126, 1205–1214 (2022).

36. Srivastava, S. K. et al. Myb overexpression overrides androgen depletion-induced cell cycle arrest and apoptosis in prostate cancer cells, and confers aggressive malignant traits: potential role in castration resistance. Carcinogenesis 33, 1149–1157 (2012).

37. Anand, S. et al. MYB sustains hypoxic survival of pancreatic cancer cells by facilitating metabolic reprogramming. EMBO Rep. 24, e55643 (2023).

38. Scholl, C. & Fröhling, S. Exploiting rare driver mutations for precision cancer medicine. Curr. Opin. Genet. Dev. 54, 1–6 (2019).

39. Vivelo, C. et al. The Landscape of US and Global Rare Tumor Research Programs: A Systematic Review. The Oncologist 29, 106–116 (2024).

40. Gupta, S. et al. A Pan-Cancer Study of Somatic *TERT* Promoter Mutations and Amplification in 30,773 Tumors Profiled by Clinical Genomic Sequencing. J. Mol. Diagn. 23, 253–263 (2021).

41. Frankish, A. et al. GENCODE reference annotation for the human and mouse genomes. Nucleic Acids Res. 47, D766–D773 (2019).

42. Grant, C. E., Bailey, T. L. & Noble, W. S. FIMO: scanning for occurrences of a given motif. Bioinformatics 27, 1017–1018 (2011).

43. Bailey, T. L., Johnson, J., Grant, C. E. & Noble, W. S. The MEME Suite. Nucleic Acids Res. 43, W39–W49 (2015).

44. Castro-Mondragon, J. A. et al. JASPAR 2022: the 9th release of the open-access database of transcription factor binding profiles. Nucleic Acids Res. 50, D165–D173 (2022).

45. Sondka, Z. et al. The COSMIC Cancer Gene Census: describing genetic dysfunction across all human cancers. Nat. Rev. Cancer 18, 696–705 (2018).

46. Ernst, J. & Kellis, M. ChromHMM: automating chromatin-state discovery and characterization. Nat. Methods 9, 215–216 (2012).

47. Hong, C., Thiele, R. & Feuerbach, L. GenomeTornadoPlot: a novel R package for CNV visualization and focality analysis. Bioinformatics 38, 2036–2038 (2022).

48. Reback, J. et al. pandas-dev/pandas: Pandas 1.0.5. Zenodo (2020) doi:10.5281/zenodo.3898987.

49. Harris, C. R. et al. Array programming with NumPy. Nature 585, 357–362 (2020).

50. Virtanen, P. et al. SciPy 1.0: fundamental algorithms for scientific computing in Python. Nat. Methods 17, 261–272 (2020).

51. Li, X. et al. 5’isomiR-183-5p|+2 elicits tumor suppressor activity in a negative feedback loop with E2F1. J. Exp. Clin. Cancer Res. CR 41, 190 (2022).

52. Alexandrov, L. B. et al. The repertoire of mutational signatures in human cancer. Nature 578, 94–101 (2020).

